# The RING-finger domain of *Arabidopsis* RMR functions as an E3 ligase essential for post-Golgi trafficking

**DOI:** 10.1101/2024.12.17.628466

**Authors:** Shuai Chen, Yonglun Zeng, Hiu Yan Wong, Yihong Chen, Lei Yang, Fang Luo, Caiji Gao, Liwen Jiang, Kam-Bo Wong

**Author notes:** To whom correspondence should be addressed. Email: Liwen Jiang; Kam-Bo Wong. These authors contributed equally to this article (Shuai Chen, Yonglun Zeng, Hiu Yan Wong).

## Abstract

Receptor-homology-transmembrane-RING-H2 (RMR) sorting receptors are essential for directing soluble cargo proteins to protein storage vacuoles in plants. These type I integral membrane proteins comprise a single transmembrane domain, an N-terminal lumenal region containing a protease-associated domain for cargo recognition, and a C-terminal cytoplasmic region (CT) with a Really-Interesting-New-Gene-H2 (RING-H2) domain. Here, we determined the crystal structure of the RING-H2 domain of *Arabidopsis* RMR isoform-1 (AtRMR1-RING), where the conserved C3H2C3 motif coordinates two Zn ions, a feature typical of RING-type E3 ligases. AtRMR1-RING was shown to interact with *Arabidopsis* E2 ubiquitin-conjugating enzyme, and exhibits E3 ligase activity in an in vitro ubiquitination assay. Biochemical analysis reveals that I234Y substitution disrupted the E2/E3 interaction and greatly reduced E3 ligase activity. Furthermore, we showed that the conserved RING-H2 domains of AtRMR isoform 2, 3 and 4 are also E3 ligases. Inactivation of E3 ligase activity by the I234Y mutation resulted in Golgi retention of AtRMR1-CT and AtRMR2. These findings suggest that the E3 ligase activity is essential for post-Golgi trafficking of RMR receptors, providing new insights into receptor-mediated protein sorting in plants.

## Introduction

Plant cells are characterized by a prominent vacuolar system, which occupies a significant portion of the cell volume (1). This system consists primarily of two types of vacuoles: lytic vacuoles and protein storage vacuoles (PSVs). Lytic vacuoles play roles in various cellular processes, including metabolite recycling, defence against infection, autophagy and programmed cell death (2). In contrast, PSVs are responsible for the accumulation of storage proteins in developing seeds. A key mechanism by which soluble cargo proteins are sorted to plant vacuoles involves the action of sorting receptors (3). Specifically, the sorting of soluble cargo proteins to vacuoles is mediated by type I transmembrane receptors, namely vacuolar sorting receptors (VSR) and receptor-homology-transmembrane-RING-H2 (RMR) proteins (4). Both receptor families share common structural features, including an N-terminal lumenal region (NT) responsible for cargo binding, and C-terminal cytoplasmic regions (CT) that are crucial for targeting and subcellular localization (5–12).

RMR contains a protease-associated (PA) domain in the NT, separated by a transmembrane domain (TMD), and a Really-Interesting-New-Gene-H2 (RING-H2) domain in the CT (Fig. 1A). The PA domain, responsible for receptor-cargo interactions, is present in both VSR and RMR (5, 7, 9, 10, 12, 13), whereas the CT of these receptors differ markedly. VSR contains a short cytosolic tail with the conserved YxxΦ and acidic dileucine-like motifs that determine its trafficking (3, 14), whereas RMR has a distinct C-terminal RING-H2 domain (11, 15) (Fig. 1A). Early studies revealed that the NT of RMR is responsible for directing the reporter protein pro-aleurain to storage vacuoles via the Golgi apparatus (6). In contrast, fusion of RMR’s NT with TMD and CT of VSR resulted in the mislocalization of the fusion product to lytic vacuoles (16). These findings suggest that RMR and VSR follow distinct trafficking pathways, with the CT of RMR playing a crucial role in the proper targeting and transport of cargo to its correct destination.

**Figure 1.**
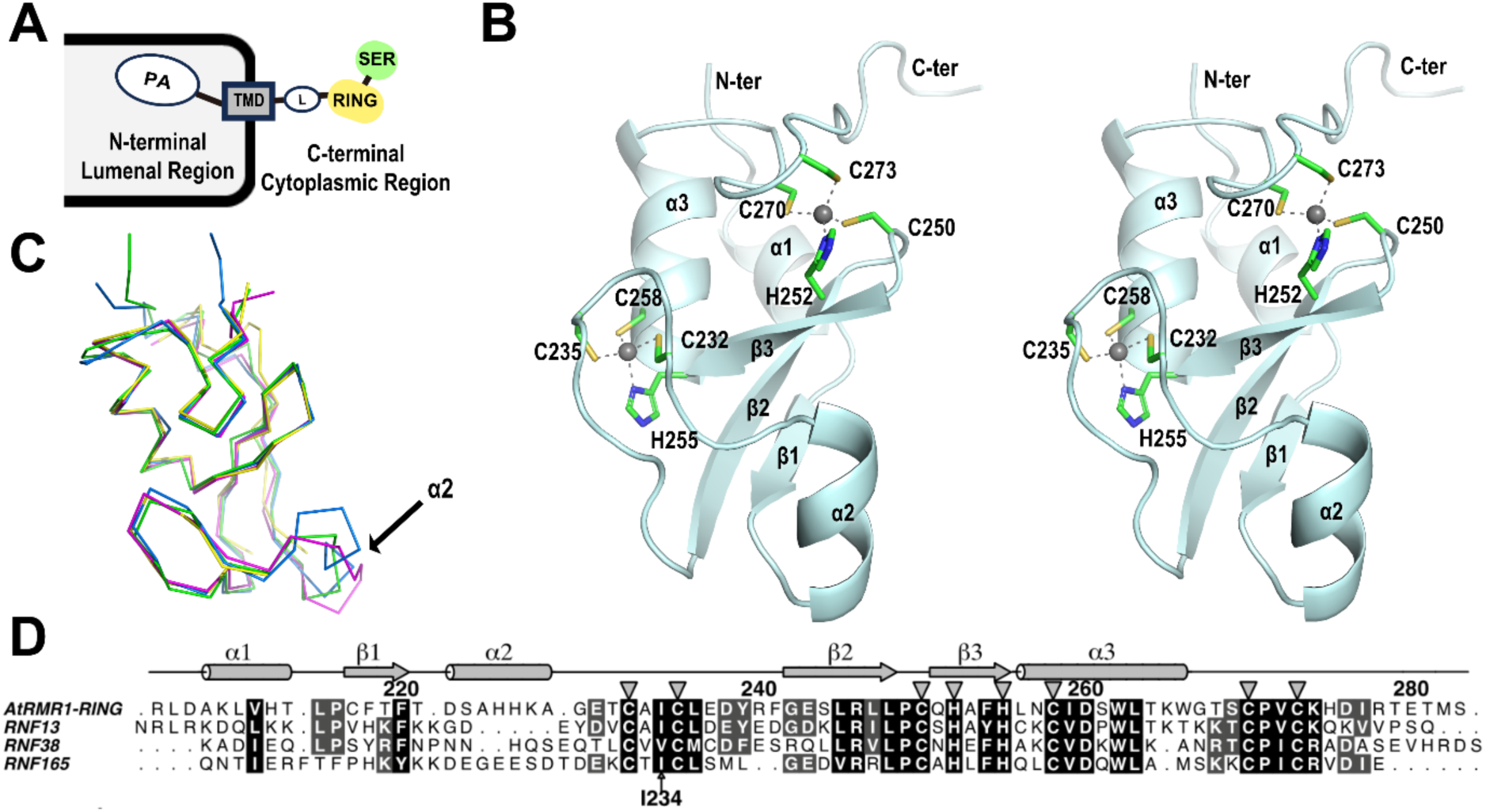
Crystal structure of AtRMR1-RING resembles an E3 ligase. (A) Domain organization of receptor-homology-transmembrane-RING-H2 (RMR) sorting receptors. RMR contains a protease-associated (PA) domain at its N-terminal lumenal region for cargo-binding. The C-terminal cytoplasmic region of RMR contains a RING-H2 (RING) domain connected to the transmembrane domain (TMD) via a Linker (L) domain, and a serine-rich (SER) domain. (B) Ribbon representations of the crystal structure of the RING-H2 domain of *Arabidopsis* RMR isoform-1 (AtRMR1-RING). The conserved C3H2C3 motif (Cys232, Cys235, Cys250, His252, His255, Cys258, Cys270 and Cys273) coordinating two Zn ions (grey spheres) is shown. Secondary structures of AtRMR1-RING are also labeled. (C) The structure of AtRMR1-RING (blue) is superimposable with the crystal structures of RNF13 (green, PDB code: 5ZBU), RNF38 (magenta, PDB code: 4V3K) and RNF165 (yellow, PDB code: 5D0M). Helix-2 of AtRMR1-RING is not conserved in other RING-H2 domains. (D) Sequence alignment of AtRMR1 with RNF13, RNF38 and RNF165. AtRMR1-RING shares 42%, 31% and 34% sequence identity to the RING-H2 domain of RNF13, RNF38 and RNF165, respectively.

Despite these insights, the biological function of the C-terminal RING-H2 domain of RMR remains poorly understood. Interestingly, this domain is also present in other mammalian E3 ligases known to be involved in protein trafficking (17). In this study, we determined the crystal structure of the RING-H2 domain of Arabidopsis RMR isoform 1 (AtRMR1-RING), revealing its structural similarity to known E3 ligases. Our *in vitro* studies further confirm that the RING-H2 domain of AtRMR1-4 functions as an E3 ligase. Additionally, we show that I234Y substitution, which disrupts E3 ligase activity, results in the retention of RMR in the Golgi apparatus of *Arabidopsis* protoplasts, highlighting the essential role of ubiquitination in protein trafficking.

## Results

### Crystal structure of AtRMR1-RING

RMR sorting receptors are type-1 transmembrane protein that adopt a PA-TMD-RING domain organization, with a protease-associated (PA) domain at the N-terminal region (NT), a transmembrane domain (TMD), and a RING-H2 domain at the C-terminal region (CT) (Fig. 1A). To better understand how the CT of RMR affects protein trafficking, we determined the crystal structure of the RING-H2 domain of AtRMR1 (AtRMR1-RING, residue 206-283) to a resolution of 2.1 Å (Fig. 1B and Table S1). There are two AtRMR1-RING in the crystallographic asymmetric unit (Fig. S1A) and each AtRMR1-RING binds two zinc ions by the C3H2C3 motif – the first zinc ion is coordinated in a tetrahedral geometry by the _232_CxxC_235_ and _252_HxxC_255_ motifs, while the second one by the _250_CxH_252_ and _270_CxxC_273_ motifs (Fig. S1B). The C3H2C3 motif is conserved in the RING-H2 domain of other known E3 ligases with the same PA-TMD-RING topology (Fig. S1C).

We further compared the crystal structure of AtRMR1-RING with other known crystal structures of RING-H2 domain from RNF13, RNF38 (18), and RNF165 (19) (Fig. 1C and 1D). The structure of AtRMR1-RING is similar to other RING-H2 domains, adopting the typical RING finger fold (20). However, helix-2 of AtRMR1-RING is not found in RNF13 and RNF38, while the corresponding residues in RNF165 are disordered.

### AtRMR1-RING is an E3 ligase

Given that the sequence and structure of AtRMR1-RING resembles those of known E3 ligases, it is likely that AtRMR1-RING is also an E3 ligase. As glutathione-S-transferase (GST) has been previously shown to serve as a substrate for E3 ligase in an auto-ubiquitination assay (21), we tested if AtRMR1-RING is an E3 ligase by mixing GST-tagged AtRMR1-RING (GST-AtRMR1-RING) with *Arabidopsis* E1 ubiquitin-activating enzyme (AtUba1, At2g30110), E2 ubiquitin-conjugating enzyme (AtUbc30, At5g56150) and ubiquitin (AtUb, At4g02890) in the presence of Mg^2+^ and ATP. Poly-ubiquitination was detected by Western blot using the FK2 antibody (ENZO Biochem), which is specific for poly-ubiquitin but not free ubiquitin (Fig. S3). In the presence of both E1 and E2 enzymes, poly-ubiquitination was detected in GST-AtRMR1-RING but not in the GST control (Fig. 2A), suggesting that AtRMR1-RING is indeed an E3 ligase.

**Figure 2.**
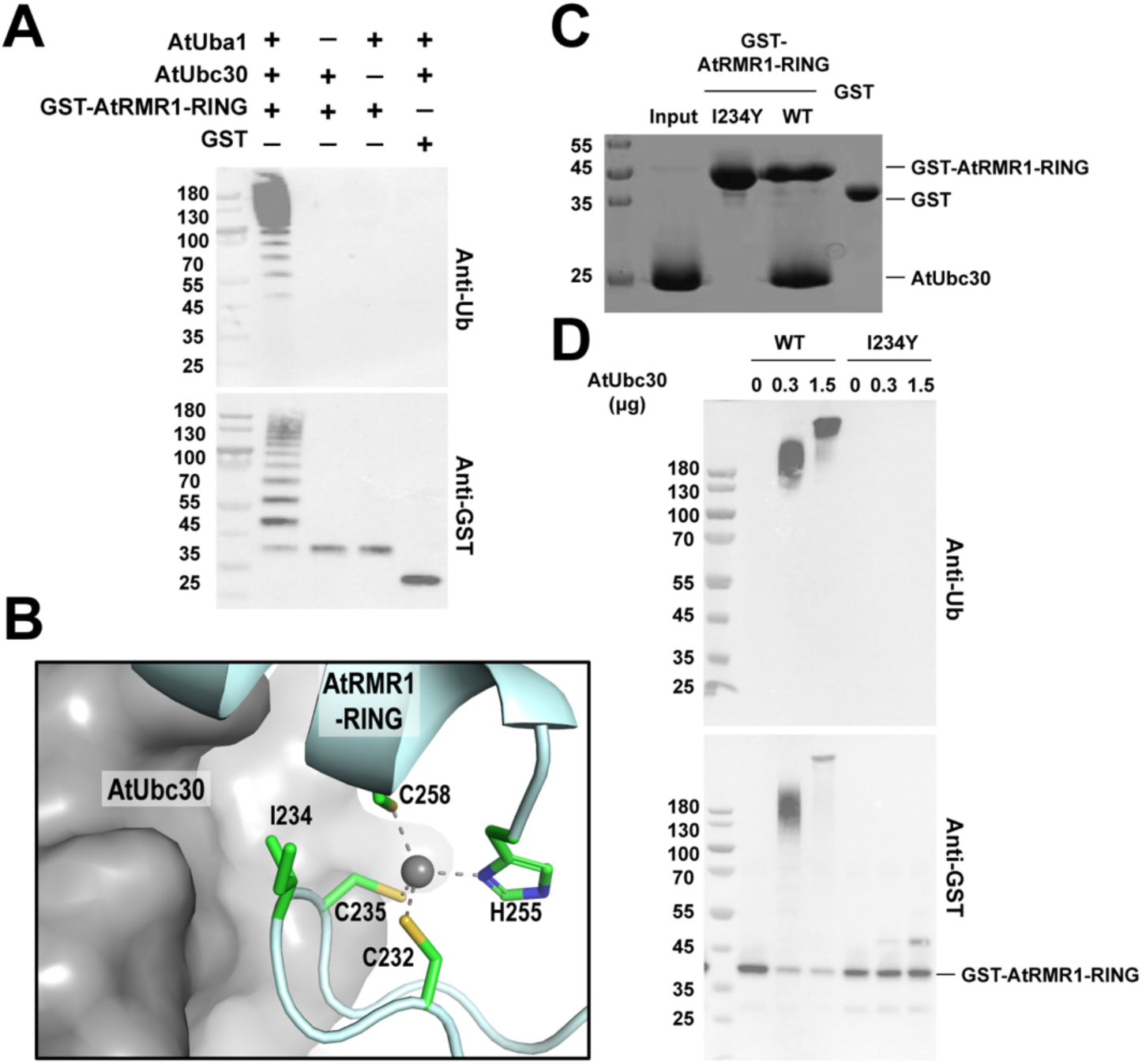
I234Y substitution disturbed the E3 ligase activity of AtRMR1-RING. (A) *In vitro* ubiquitination assay showing AtRMR1-RING is an E3 ligase. 1 µg GST-tagged AtRMR1-RING or GST were mixed with 0.1 µg Arabidopsis E1 ubiquitin-activating enzyme (AtUba1), 0.3 µg E2 ubiquitin-conjugating enzyme (AtUbc30), 2.5 µg ubiquitin (AtUb) in the presence of Mg^2+^ and ATP at 30 °C, 30 min and analyzed by Western blotting using ubiquitin (FK2, Enzo Life Sciences) and GST (Abcam) antibodies. (B) How AtRMR1-RING (cyan) interacts with AtUbc30 (grey) was predicted by homology modelling based on the crystal structures of the human RNF38/UbcH5B/Ubiquitin complex (PDB code: 4V3K). The conserved Ile234 of AtRMR1-RING is located at the interface between AtRMR1-RING and AtUbc30. (C) I234Y substitution disturbed the interaction between AtRMR1-RING and AtUbc30. 50 µM purified AtUbc30 (Input) was mixed with 50 µM wild-type, I234Y variant of GST-AtRMR1-RING or GST in the presence of GSTrap Sepharose (GE Healthcare). After extensive washing, bound proteins were eluted with 20 mM reduced glutathione and analyzed by 15% SDS-PAGE with Coomassie blue staining. AtUbc30 was co-eluted with wild-type GST-AtRMR1-RING but not with the I234Y variant nor the GST control. (D) I234Y substitution greatly reduced the E3 ligase activity of AtRMR1-RING. Wild-type or I234Y variant of GST-AtRMR1-RING (1 µg) was incubated with AtUba1 (0.1 µg), AtUbc30 (0 µg, 0.3 µg or 1.5 µg) and AtUb (2.5 µg) in the presence of Mg^2+^ and ATP at 30 °C, 30 min. Poly-ubiquitination was detected by Western blotting using ubiquitin and GST antibodies. Poly-ubiquitination was evident in wild-type GST-AtRMR1-RING, but not in the I234Y variant, suggesting that I234Y substitution greatly reduced the E3 ligase activity.

### I234Y substitution greatly reduced the E3 ligase activity of AtRMR1-RING

To further understand how AtRMR1-RING interacts with its E2 ubiquitin-conjugating enzyme, we modelled the complex structure of AtRMR1-RING/AtUbc30 based on the complex structure of RNF38/UbcH5B/ubiquitin (18) (Fig. S2). The model suggests that Ile234 of AtRMR1 undergoes hydrophobic interactions with the E2 ubiquitin-conjugating enzyme (Fig. 2B). Interestingly, Ile234 is highly conserved and this residue has been shown to be crucial for the activity of other RING E3 ligases (22, 23).

To test the role of Ile234 in AtRMR1, we created an I234Y variant of AtRMR1-RING. First, we performed analytical gel filtration on purified protein samples to check if the I234Y substitution affects proper folding of AtRMR1-RING. The elution volume of the I234Y variant was ∼14.4 ml (Fig. S4), which is similar to that of the wild-type AtRMR1-RING. This observation suggests that the I234Y variant and the wild-type AtRMR1-RING have similar hydrodynamic radii and are properly folded. In contrast, when wild-type AtRMR1-RING and the I234Y variant were denatured by addition of 8M urea to the protein samples and the running buffer, the elution volumes of wild-type AtRMR1-RING and the I234Y variant were shifted to ∼12.7 and ∼12.6 ml, respectively (Fig. S4). Next, we tested the ability of the I234Y variant to form an E2/E3 complex via a GST pull-down assay (Fig. 2C). Substitution of a bulky tyrosine residue was expected to break the E2/E3 interaction by creating steric clashes in the interface (Fig. 2B). Our results showed that while wild-type GST-AtRMR1-RING can interact with AtUbc30, the I234Y substitution abolished the interaction between AtRMR1-RING and AtUbc30 (Fig. 2C).

As RING-type E3 ligases promote catalysis by forming an E2/E3 complex so that the substrate is juxtaposed to the E2-ubiquitin thioester (24), we expected that the I234Y substitution should inactivate the E3 ligase activity of AtRMR1-RING. To test this hypothesis, we performed the *in vitro* ubiquitination assay and showed that I234Y substitution did indeed greatly reduced the E3 ligase activity of AtRMR1-RING (Fig. 2D).

### E3 ligase activity of AtRMR1 is essential for its protein trafficking

To further investigate the *bona fide* function of AtRMR1-RING as an E3 ligase and its impacts on protein trafficking, we performed an *in vivo* analysis using our established *Arabidopsis* protoplast transient expression system (25). It has been shown that the CT is involved in targeting individual VSRs and RMRs to their final destinations (6–8), we thus used similar strategies to study the localization of AtRMR1 as well as the I234Y mutant. We used ManI-mRFP and mRFP-SYP61 as markers for the Golgi apparatus and TGN, respectively (Fig. S5). GFP-TMD-AtRMR1-CT was co-transfected with ManI-mRFP or mRFP-SYP61 (Figs. 3A, 3B, S6 and S7). GFP-TMD-AtRMR1-CT exhibited partial colocalization with both the Golgi marker and *trans*-Golgi-network (TGN) marker. Colocalization of AtRMR1-CT with the markers was quantified by the Pearson correlation analysis using the Costes’ automatic threshold method (26). The Pearson correlation coefficient (r_p_) for the colocalization of the I234Y mutant with ManI-mRFP (r_p_ = 0.90 ± 0.01) was significantly higher than that of the wild-type (r_p_ = 0.54 ± 0.03) (Fig. 3C), suggesting that the I234Y substitution increases the colocalization of AtRMR1-CT with the Golgi marker. On the other hand, the correlation coefficient for the colocalization of I234Y mutant with mRFP-SYP61 (r_p_ = 0.27 ± 0.14) was not significantly different from that of the wild-type (r_p_ = 0.52 ± 0.09) (Fig. 3D). Our results suggest that the I234Y substitution leads to an increased retention of the AtRMR1-CT in the Golgi apparatus.

**Figure 3.**
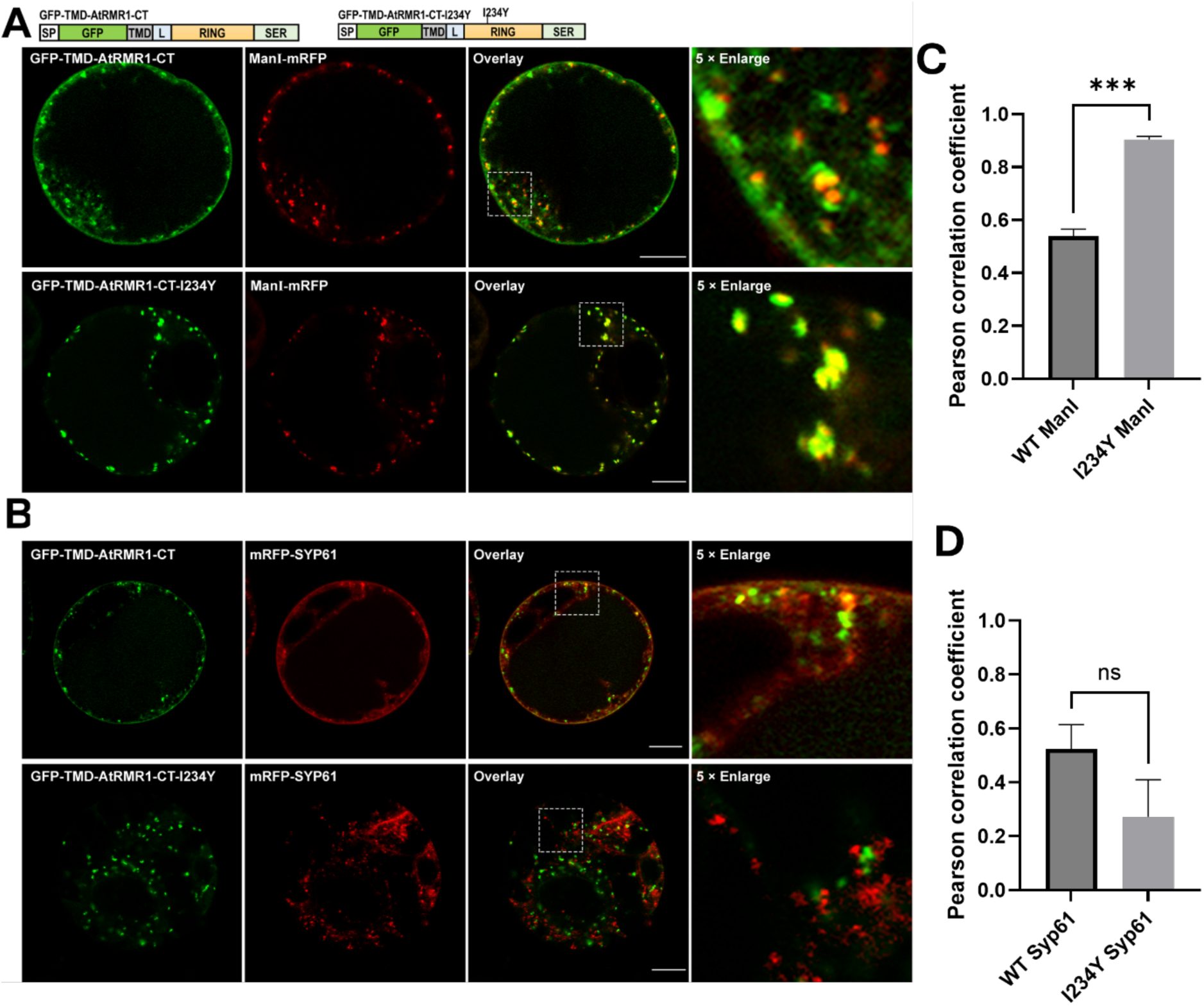
I234Y substitution resulted in Golgi retention of GFP-TMD-AtRMR1-CT in *Arabidopsis* protoplasts. *Arabidopsis* protoplasts were co-transfected with wild-type or I234Y mutant of GFP-TMD-AtRMR1-CT and the (A) Golgi (ManI-mRFP) or (B) TGN (mRFP-SYP61) markers before confocal imaging of transfected cells. (Scale bar, 10 μm). Colocalization of GFP-TMD-AtRMR1-CT and the (C) Golgi or (D) TGN markers in the enlarged images were quantified by Pearson colocalization coefficient (r_P_) using the Costes’ automatic threshold. Data represent mean ± SEM (n=3). The r_P_ values for the I234Y mutant and ManI-mRFP was significantly higher than that of the wild-type. (P < 0.001, two-tailed t test).

### The RING domains of AtRMR2-4 are also E3 ligases

Given that AtRMR1-RING functions as an E3 ligase and the RING-H2 domain is conserved across AtRMR homologs (Fig. 4A), we hypothesized that they might exhibit similar E3 ligase activity. To address this hypothesis, we tagged GST to the N-termini of the RING-H2 domain of AtRMR2-4 and tested their E3 ligase activity (Fig. 4B-D). In the presence of both E1 and E2 enzymes, polyubiquitination was detected in GST-AtRMR2-4-RING, but not in the GST control. These observations suggest that the RING-H2 domain of AtRMR2-4 are E3 ligases.

**Figure 4.**
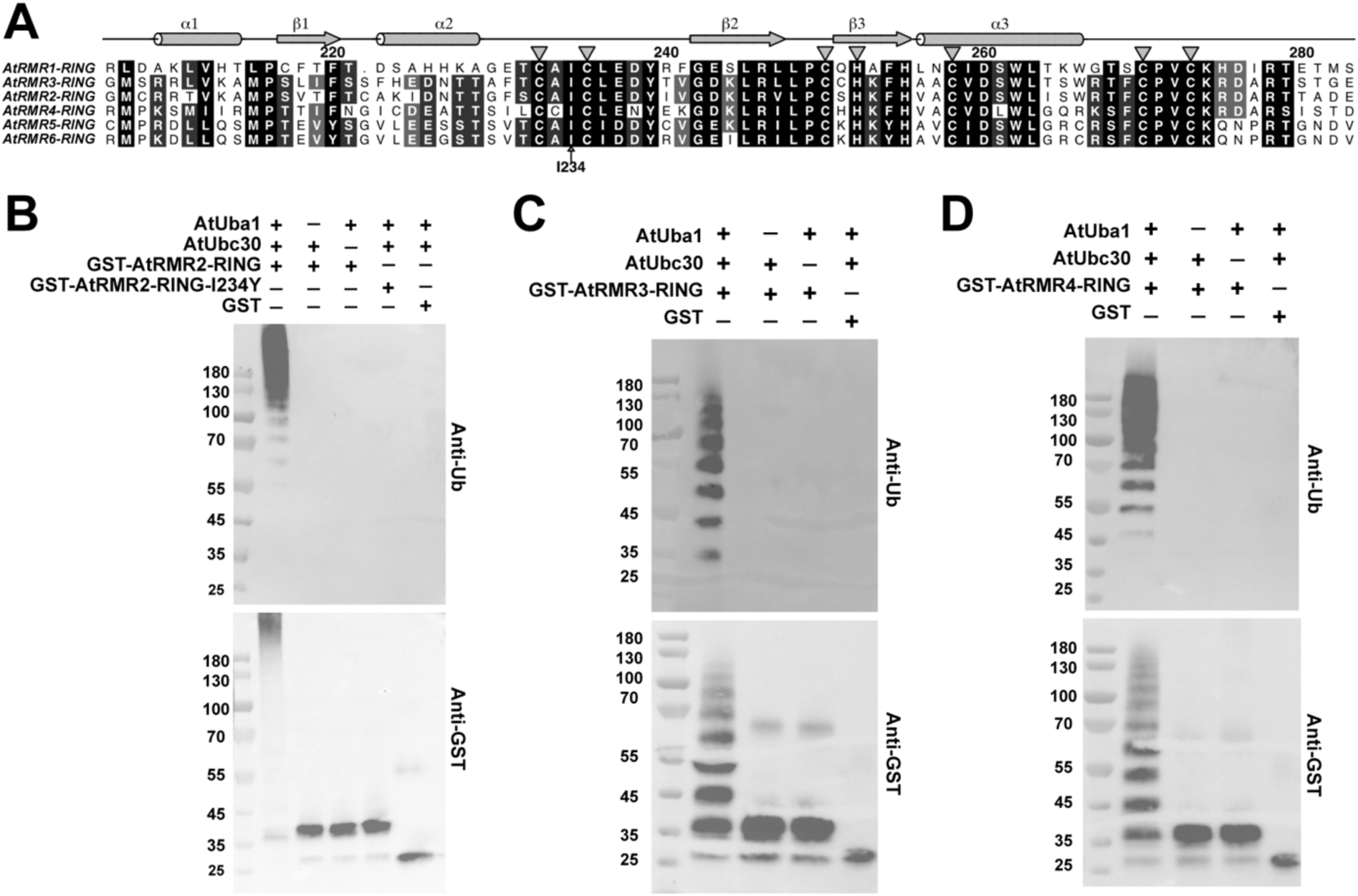
The RING domains of AtRMR2-4 are E3 ligases. (A) Sequence alignment of the RING domains of AtRMR1-6, showing conservation of the C3H2C3 motif (indicated by triangles) and Ile234 in AtRMR. (B) I234Y substitution disrupted E3 ligase activity of AtRMR2-RING. In the E3 ligase activity assay, GST, GST-tagged AtRMR2-RING, or its I234Y variant were mixed with Arabidopsis E1 ubiquitin-activating enzyme (AtUba1), E2 ubiquitin-conjugating enzyme (AtUbc30), and ubiquitin in the presence of Mg^2+^ and ATP at 30°C, 30 min and analyzed by Western blotting using ubiquitin (FK2, Enzo Life Sciences) and GST (Abcam) antibodies. In the presence of both E1 and E2, poly-ubiquitination was evident in wild-type GST-AtRMR2-RING but not in the I234Y variant. (C & D) The E3 ligase assay was performed for GST-tagged AtRMR3-RING or AtRMR4-RING. In the presence of both E1 and E2, poly-ubiquitination was observed for GST-AtRMR3-RING and GST-AtRMR4-RING.

### I234Y substitution led to Golgi retention of AtRMR2

Similar to AtRMR1, AtRMR2 also contains the conserved Ile234 residue in its RING-H2 domain (Fig. 4A). We also show that the I234Y substitution abolished the E3 ligase activity of AtRMR2-RING (Fig. 4B). Next, we investigated if the I234Y mutation affects the trafficking of AtRMR2 by co-transfecting GFP-AtRMR2 with the Golgi or TGN markers in *Arabidopsis* protoplasts (Figs. 5A, 5B, S8 and S9) and colocalization was quantified by the Pearson correlation coefficient (Figs. 5C and 5D). Consistent with previous results (11), GFP-AtRMR2 partially colocalized with the Golgi and TGN markers. The correlation coefficient for the colocalization of the I234Y mutant of AtRMR2 with ManI-mRFP (r_p_ = 0.63 ± 0.01) was significantly higher than that of the wild-type (r_p_ = 0.34 ± 0.04) (Fig. 5C), suggesting that the I234Y substitution increased the colocalization with the Golgi marker. On the other hand, the correlation coefficient for the colocalization of the I234Y mutant with mRFP-SYP61 (r_p_ = 0.39 ± 0.11) was not significantly different from that of the wild-type (r_p_ = 0.59 ± 0.09) (Fig. 5D). These findings suggest that inactivation of the E3 ligase activity by the I234Y substitution leads to retention of both GFP-TMD-AtRMR1-CT and GFP-AtRMR2 in the Golgi apparatus (Fig. 3 and 5), further supporting the conclusion that E3 ligase activity of the CT of RMR receptors is essential for their proper post-Golgi trafficking.

**Figure 5.**
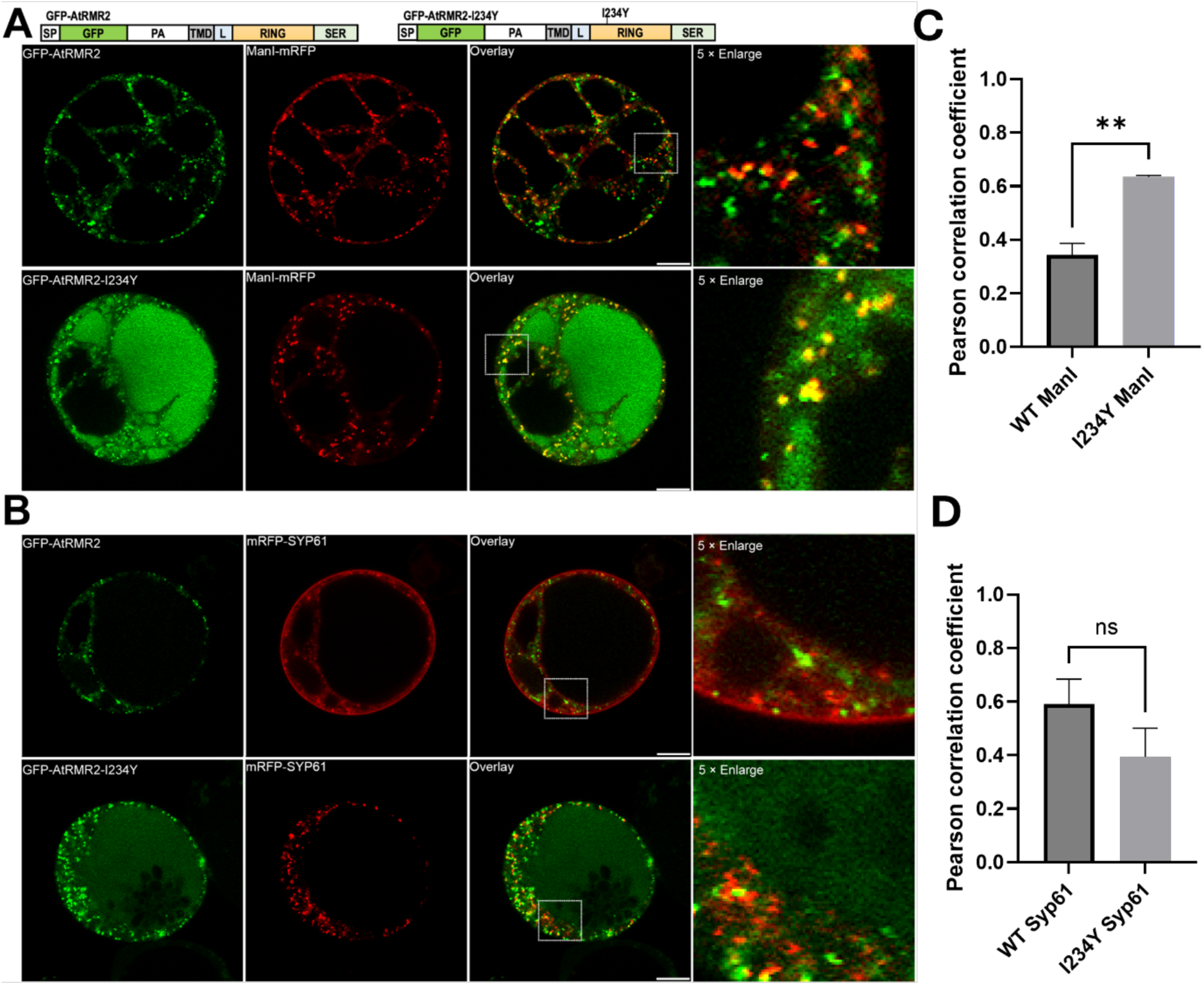
I234Y substitution resulted in Golgi retention of GFP-AtRMR2 in *Arabidopsis* protoplasts. *Arabidopsis* protoplasts were co-transfected with wild-type or I234Y mutant of GFP-AtRMR2 and the (A) Golgi (ManI-mRFP) or (B) TGN (mRFP-SYP61) markers before confocal imaging of transfected cells. (Scale bar, 10 μm). Colocalization of GFP-AtRMR2 and the (C) Golgi or (D) TGN markers in the enlarged images were quantified by Pearson colocalization coefficient (r_P_) using the Costes’ automatic threshold. Data represent mean ± SEM (n=3). The r_P_ values for the I234Y mutant and ManI-mRFP was significantly higher than that of the wild-type. (P < 0.01, two-tailed t test).

## Discussion

One prominent feature of RMR is the presence of a conserved RING-H2 domain in the CT. While the CT of RMR has been implicated in protein targeting (6, 7, 9, 11), the underlying mechanism remains elusive. Due to its sequence similarity to other well-characterized RING-type E3 ligases (Fig. S1), the role of RMR proteins as E3 ligases has long been hypothesized (15) but not demonstrated experimentally. Here, we present the crystal structure of the RING-H2 domain of AtRMR1 (Fig. 1) and demonstrated that the RING-H2 domain of RMR1-4 are E3 ligases in a biochemical assay (Figs. 2 and 4).

We further showed that the I234Y mutation disturbing the E3 ligase activity resulted in Golgi retention of GFP-TMD-AtRMR1-RING and GFP-AtRMR2 in *Arabidopsis* protoplasts (Fig. 3 and 5). Our observation is consistent with a pioneer study of Park and co-workers, who showed that truncation of the CT of AtRMR1 resulted in its colocalization with the Golgi apparatus and blocked its trafficking to dark intrinsic protein (DIP) organelles in *Arabidopsis* protoplasts (7). It is tempting to speculate that the E3 ligase activity of RMR provides a novel mechanism to regulate its post-Golgi trafficking such that inactivating the E3 ligase activity by I234Y substitution resulted in aberrant accumulation of RMR in the Golgi apparatus.

It has been well established that ubiquitination plays an essential role in the endocytic pathway and in protein degradation to lysosomes/vacuoles (27–30). In yeast and mammalian cells, GGA (Golgi-localized, γ-adaptin ear-containing, ADP ribosylation factor-binding protein) serves as an Endosomal-Sorting-Complex-Required-for-Transport (ESCRT)-0-like component to convey ubiquitinated cargos to the lysosomal degradation (30–32). Although GGA is absent in plants, its role is likely replaced by a related TOL (Tom-1 Like) protein, which also contains the conserved VHS and GAT domains for cargo recognition (28). For example, *Arabidopsis* TOL2 and TOL6 serve as receptors for ubiquitinated proteins and are involved in the vacuolar degradation of plasma membrane proteins via the ESCRT pathway (33, 34). In a recent study, Xu and co-workers showed that recognition of the ubiquitination on Snc1, a SNARE protein, by the WD40 domain of COPI coatomers is essential for the recycling of Snc1 in yeast (35). They argued that poly-ubiquitination provides a COPI-dependent sorting signal that redirects cargos from the endocytic pathway to the TGN and is independent to the known COPI-mediated retrograde transport of cargos from the Golgi apparatus to the ER (35). A similar scenario was reported for RNF11 (ring finger protein 11), a RING-H2 E3 ligase that drives the ubiquitination of GGA3 and governs RNF11 sorting at the TGN, in mammalian cells (36). Accordingly, a lysine-less mutant of RNF11, which cannot be ubiquitinated, was found to accumulate in the Golgi (37).

In plants, there are two types of vacuolar sorting receptors, namely VSR and RMR. Both VSR and RMR are type-1 transmembrane proteins, which contain a PA domain in the N-terminal lumenal domain that can recognize the sequence-specific information known as vacuolar sorting determinant of soluble cargo proteins and sort them to the vacuoles (5, 7, 9, 10, 12, 13). The C-terminal cytoplasmic region, responsible for targeting and subcellular localization, are different in VSR and RMR. VSR has a short cytoplasmic tail containing a YxxΦ and acidic dileucine-like motifs (3, 14) while RMR contains a RING-H2 domain (11, 15) in their C-terminal region. VSR travels through endoplasmic reticulum – Golgi – TGN to a prevacuolar compartment (PVC) (38). After releasing its cargos, VSR can be recycled back to the TGN or the Golgi apparatus mediated by the retromer complex or by a clathrin-dependent mechanism (39, 40).

The trafficking pathway of RMR is less-well understood. In developing seeds, RMR is targeted to the protein storage vacuoles (PSV) (6, 15, 41). The presence of complex glycans in RMR strongly suggests its trafficking through the Golgi apparatus (6). Our results suggest that the retention of the I234Y variant of RMR in the Golgi apparatus is due to blockage of post-Golgi pathways that are dependent on the E3 ligase activity. RMR was also found in dense vesicles (DV) and in the storage PVC or DIP organelles (7, 42, 43). Unlike VSR, it has been proposed that RMR is involved in the aggregation of storage proteins from the Golgi into dense vesicles, which are then internalized into storage PVC during their passage to the PSV (15). Given that ubiquitin was also found in the PSV (15) and the RING-H2 domain is only found in RMR but not in VSR, it would be tempting to speculate that trafficking from the Golgi apparatus to DV or DIP organelles is dependent on E3 ligase activity. Blockage of this pathway by I234Y substitution may lead to the aberrant accumulation of RMR in the Golgi apparatus. We also noticed that the presence of both wild-type AtRMR2 and the I234Y variant in the vacuoles (Figs. 5, S8 and S9). These observations may suggest that AtRMR2, like VSR, can also be sorted to the vacuoles via an E3-ligase independent pathway.

It is unclear if cargo-binding affects trafficking of RMR. Since the PA domain of RMR responsible for cargo binding is facing the lumen but the RING-H2 domain is facing the cytoplasm, it is unlikely that RMR can ubiquitinate its cargo proteins. However, it is interesting to note that phaseolin, a seed storage protein that is known to bind RMR, adopts a trimeric quaternary structures (44). Conserved lysine residues are found within the RING-H2 domain of AtRMR1-6. Binding of phaseolin could bring together multiple RING-H2 domains of RMR in close proximity, which may promote auto-ubiquitination or ubiquitination of its substrates. It would be interesting to identify the substrate of RMR in the future to provide a better understanding of how the E3 ligase activity of RMR affects its post-Golgi trafficking.

## Experimental procedures

### Plasmid construction

To create vectors for protein expression in *E. coli*, codon-optimized coding sequences of AtUba1 (72-1080), AtUbc30, AtUb were ordered from GenScript (http://www.genscript.com). AtRMR1-RING (206-283), AtRMR2-RING (207-283), AtRMR3-RING (207-282), AtRMR4-RING (209-284) were PCR amplified and all cloned into the pGEX-6p-1 vector (for GST-tagged fusion proteins) as listed in the Table S2 and S3. To create GFP-tagged constructs for transient expression in *Arabidopsis* protoplasts, coding sequences of AtRMR1-CT (160-310) and AtRMR2 (22-448) were cloned into pBI221 to create GFP-TMD-AtRMR1-CT and GFP-AtRMR2, respectively (Table S2 and S3). I234Y mutation was introduced using Q5 site-directed mutagenesis kit (New England BioLabs) with primers listed in Table S3.

### Protein Expression and Purification

Expression vectors pGEX-6p-1-AtRMR1-4-RING, pGEX-6p-1AtUba1, pGEX-6p-1-AtUbc30 or pGEX-6p-1-AtUb were transformed into *E. coli* SoluBL21 cell (Genlantis). Protein expression was induced by 0.4 mM isopropyl beta-D-1-thiogalactopyranoside at OD_600_ 0.8. After incubation at 16 ℃ for 18 h, the cells were harvested by centrifugation at 14,160g for 12 min. The cell pellet was resuspended in the 30 mM Tris buffer pH 7.6, 500 mM NaCl, 5 μM ZnCl_2_ and 5 mM dithiothreitol (DTT) and sonicated on ice. After centrifugation at 29,000×g for 36 min, the supernatant was filtered and then incubated with GSTrap Sepharose (GE Healthcare) for 2 h at 4 ℃. After extensive washing with 30 mM Tris, pH 7.6, 150 mM NaCl and 5 mM DTT, GST-tagged PreScission protease (GE Healthcare) was added to remove the GST tag by incubation at 4 ℃ overnight. The target proteins were then collected in the unbound fractions, and further purified by gel filtration using a HiLoad 26/60 Superdex 75 column (GE Healthcare) pre-equilibrated with 30 mM Tris, pH 7.5, 150 mM NaCl, 5 μM ZnCl_2_ and 2 mM DTT.

### Crystallization and Structure Determination

Crystals of AtRMR1-RING were obtained using hanging-drop-vapour-diffusion by mixing 1 μl protein solution (15 mg/ml in 30 mM Tris, pH 7.5, 150 mM NaCl, 5 μM ZnCl_2_ and 2 mM DTT) with 1 μl reservoir solution containing 0.2 M ammonium acetate, 0.1 M Bis-Tris pH 5.5, 20% w/v polyethylene glycol 3,350 at 16 ℃. Cryoprotection was achieved with 15% glycerol. Diffraction data were collected using an in-house rotating anode X-ray generator (Rigaku FR-E+), and processed with the XDS package (45) and the CCP4 program suite (46). The phase was solved by molecular replacement using the program MOLREP (47) with the crystal structure of RNF38 (PDB:4V3K) as the search template. Models were built interactively using the program COOT (48) and refined using the program PHENIX (49). All molecular graphics in the manuscript were generated with PyMOL (http://www.pymol.org).

### Modelling of AtRMR1-RING/AtUbc30 complex structure

The structure of AtUbc30 was predicted by homology modeling using the crystal structure of the human RNF38/UbcH5B complex (PDB: 4V3K) as the template. Sequences of AtUbc30 and UbcH5B were aligned using the program ClustalW (50) and edited using the program CHIMERA (51). Structure models of AtRMR1-RING in complex with AtUbc30 were generated using the program MODELLER as described (52, 53).

### Pull down assay

50-60 μM purified AtUbc30 was incubated with equimolar of GST, wild-type or I234Y variant of GST-AtRMR1-RING in the presence of GSTrap Sepharose (GE Healthcare) in a binding buffer (30 mM Tris, pH 7.5, 150 mM NaCl and 2 mM DTT) for 4 h at 4 ℃. After extensive washing with the binding buffer, bound proteins were eluted with 20 mM reduced glutathione in the binding buffer.

### Ubiquitination assay

Ubiquitination assay reaction was performed in a 30 µl reaction mixture by incubating 1 µg recombinant GST-AtRMR1-RING (as in Fig. 2), GST-AtRMR2-RING (as in Fig. 4B), GST-AtRMR3-RING (as in Fig. 4C) or GST-AtRMR4-RING (as in Fig. 4D) with 0.1 µg AtUba1, 0.3 µg AtUbc30, 2.5 µg AtUb in a reaction buffer (30 mM Tris-HCl, pH 7.5, 2 mM ATP, 5 mM MgCl_2_, 2 mM DTT and 0.5 U inorganic pyrophosphatase (Sigma)). The reaction mixture was incubated at 30 ℃ for 30 min, and then analyzed by SDS-PAGE using 4–12% gradient gel (Bolt Bis-Tris Plus Gel, Thermo Fisher Scientific) and detected by Western blot using ubiquitin (FK2, ENZO), GST (Abcam) or AtRMR1-RING antibodies.

### Antibody Validation

Poly-ubiquitin (cat. no. 60-0109-100) and free ubiquitin (cat. no. 60-0200-050) proteins were obtained from Ubiquigent Ltd. The proteins were resolved on a 16% Tricine-SDS-PAGE (54) and detected by Instant-Bands (EZBiolab) staining and Western blot using the ubiquitin antibody (FK2, ENZO Biochem).

### Analytical Gel filtration

100 µL 0.3 mg/ml wild-type (A) or I234Y variant (B) of AtRMR1-RING were loaded to a Superdex 75 Increase 10/300 GL column (GE Healthcare) pre-equilibrated with 0M in a running buffer (30 mM Tris, pH 7.5, 150 mM NaCl and 2 mM DTT). To denature the proteins, 8M urea was added to the protein samples before loading to the gel-filtration column pre-equilibrated with 8M urea in the running buffer.

### Protoplast transient expression assay

GFP-tagged AtRMR1-CT or AtRMR2 were co-transfected with Golgi (ManI-mRFP) or TGN (mRFP-SYP61) markers in *Arabidopsis* PSBD protoplasts were performed as described previously (8). Transfected protoplasts were incubated for 12–16 h before confocal imaging using Leica SP8. Colocalization was quantified by the Pearson coefficient of fluorescent signals using the Costes’ automatic threshold (26) implemented in the JACoP plug-in of ImageJ (55). Results from three independent transfections were analyzed.

### Accession numbers

The crystal structure of AtRMR1-RING was deposited in the Protein Data Bank (PDB codes: 7BVW). *Arabidopsis* sequences used in this study can be found with the following accession numbers: RMR1, At5g66160; RMR2, At1g71980; RMR3, At1g22670; RMR4, At4g09560; Uba1, At2g30110; Ubc30, At5g56150; ubiquitin, At4g02890.

## Author contributions

**S. C.** & **Y. L**.: writing – original draft, review & editing; conceptualization, visualization, resources, investigation, formal analysis. **H. Y. W.:** conceptualization, visualization, investigation, validation, formal analysis, writing – review & editing. **Y. C., L. Y., F. L. and C. J. G.:** investigation. **L. J.,** & **K. B. W.:** supervision, project administration, resources, conceptualization, visualization, funding acquisition, writing – original draft, review & editing.

## Acknowledgements

This study was supported by grants from the Research Grants Council of Hong Kong SAR (CUHK14151416, CUHK14118122, C4002-17GF, C4041-18EF, AoE/M-05/12, AoE/M403/16, C4012-16E, C4033-19E, and C2003-22W), NSFC (91854201 and 31670179) and Direct Grants from the Chinese University of Hong Kong.

