## Supplemental Figures and tables for "The RING-finger domain of *Arabidopsis* RMR functions as an E3 ligase essential for post-Golgi trafficking"

* To whom correspondence should be addressed.

Email:

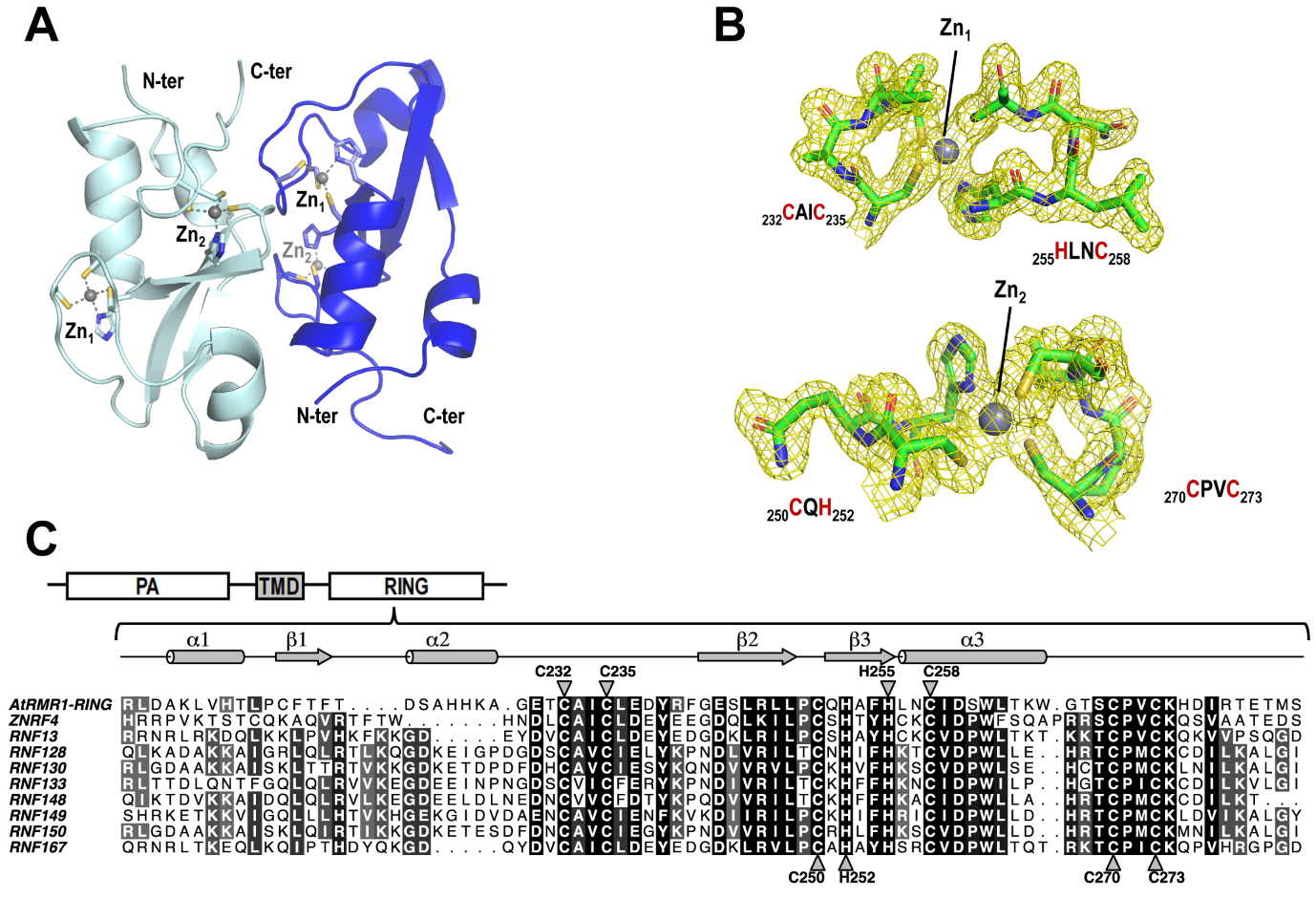

**Figure S1. Coordination of two zinc ions by the C3H2C3 motif in AtRMR1-RING.** (A) The crystallographic asymmetric unit contains two protomers (cyan and blue) of AtRMR1-RING, each binding two zinc ions. (B) Sigma-weighted 2Fo-Fc electron density map (contoured at 1σ) highlighting the two zinc-binding sites in AtRMR1-RING. Zn1 is coordinated by Cys232, Cys235, His255, and Cys238, while Zn2 is coordinated by Cys250, His252, Cys270, and Cys273. These residues form the C3H2C3 motif. (C) The C3H2C3 motif is conserved in other proteins with the PA-TMD-RING domain organization. RMR contains a protease-associated domain at its N-terminal region and a RING-H2 domain at its C-terminal region. The sequence of AtRMR1-RING is aligned with the RING-H2 domain of other known E3 ligases (ZNRF4, RNF13, RNF128 RNF130, RNF133, RNF148, RNF149, RNF150, RNF167) with this PA-TMD-RING topology. Residues involved in binding Zn1 are indicated by inverted triangles and those in binding Zn2 are indicated by triangles. Residues are numbered according to the sequence of AtRMR1. The secondary structure elements of the resolved AtRMR1-RING structure are shown above the sequence alignment.

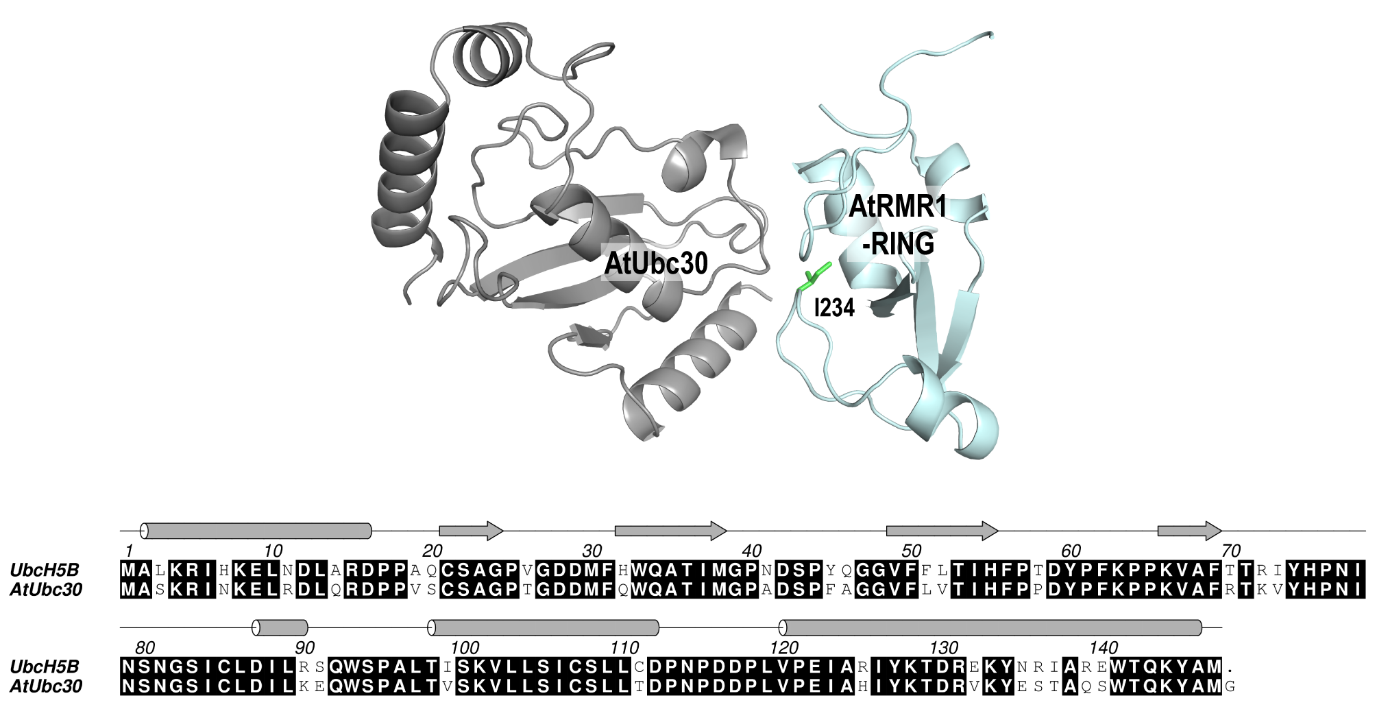

**Figure S2. Modelling of the complex structure of AtRMR1-RING/AtUbc30.** The structure of AtUbc30 is predicted using homology modelling with the crystal structure of human UbcH5B as a template. The crystal structure of AtRMR1-RING and the predicted structure of AtUbc30 are juxtaposed according to the crystal structure of RNF38-UbcH5B-Ubiqutin complex (PDB code: 4V3K). As shown in the sequence alignment, AtUbc30 shares 79% sequence identity to UbcH5B.

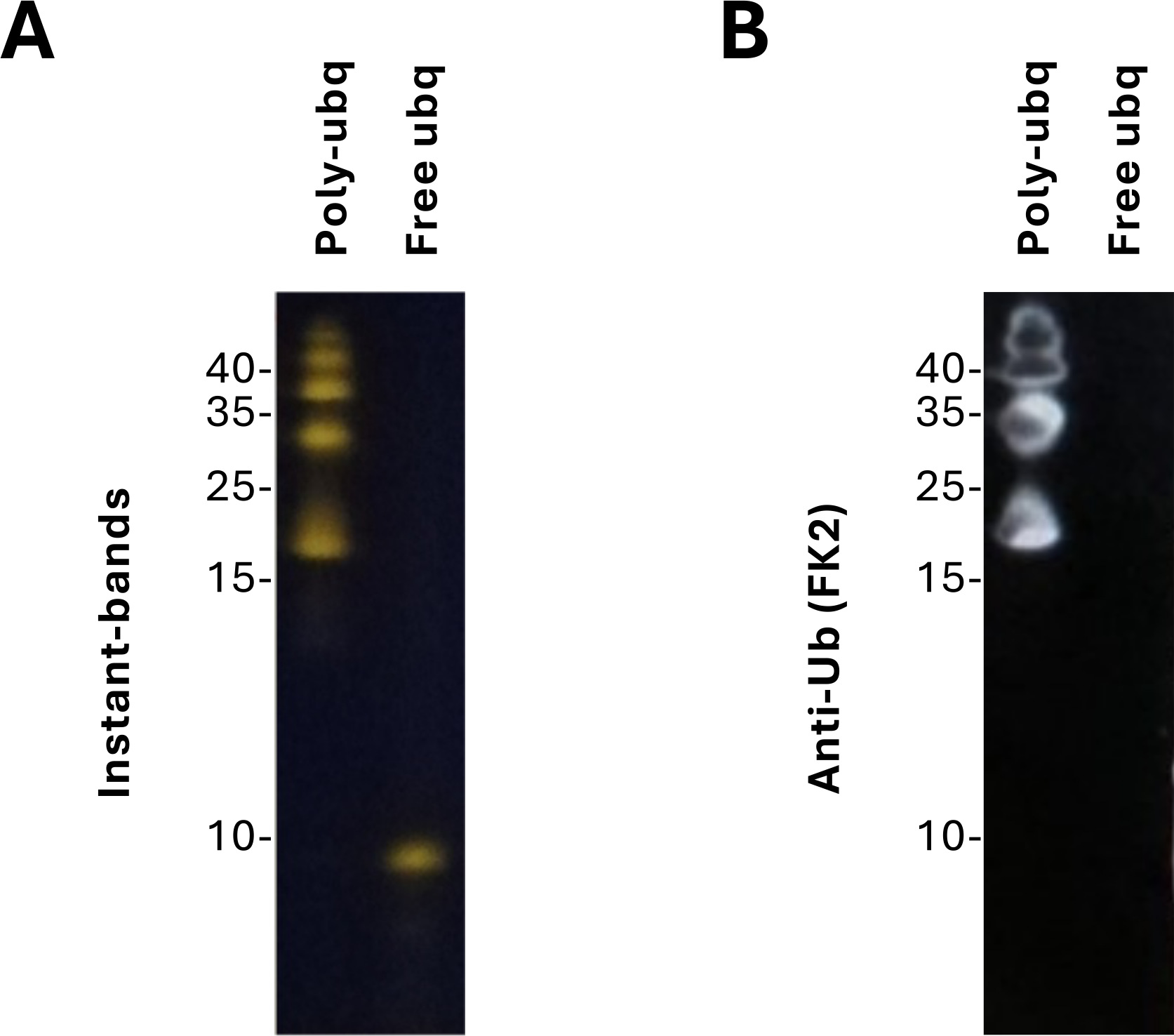

**Figure S3. The FK2 ubiquitin antibody detects poly-ubiquitin but not free ubiquitin.** Poly-ubiquitin and free ubiquitin were resolved on a 16% Tricine-SDS-PAGE. They were detected by (A) Instant-Bands (EZBiolab) that stains proteins fluorescently and (B) Western blot using the FK2 ubiquitin antibody (ENZO Biochem). We showed that the FK2 antibody detected only the poly-ubiquitin but not the free ubiquitin.

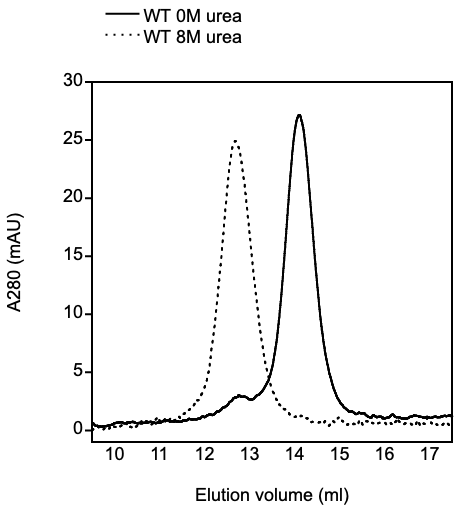

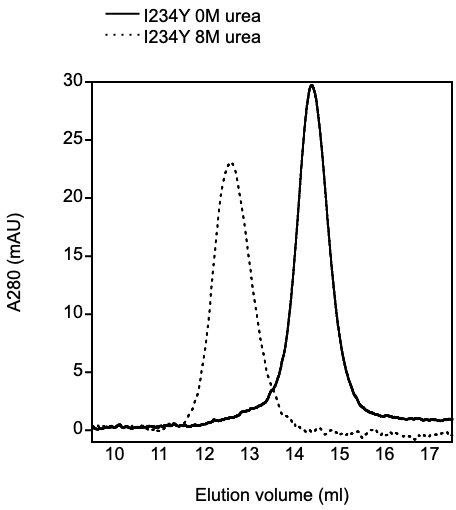

**Figure S4. Gel filtration analysis showed that the I234Y variant of AtRMR1-RING is folded.** 100 µL 0.3 mg/ml wild-type (A) or I234Y variant (B) of AtRMR1-RING were loaded to a Superdex 75 Increase 10/300 GL column (GE Healthcare) pre-equilibrated with 0M or 8M urea in a running buffer (30 mM Tris, pH 7.5, 150 mM NaCl and 2 mM DTT). The elution volume for I234Y variant was 14.4 ml, which is similar to that for WT, suggesting the I234Y variant was folded. When 8M urea was added to the protein samples and the running buffer, the elution volumes of WT and I234Y variant (dotted lines) were shifted to 12.7 and 12.6 ml, respectively.

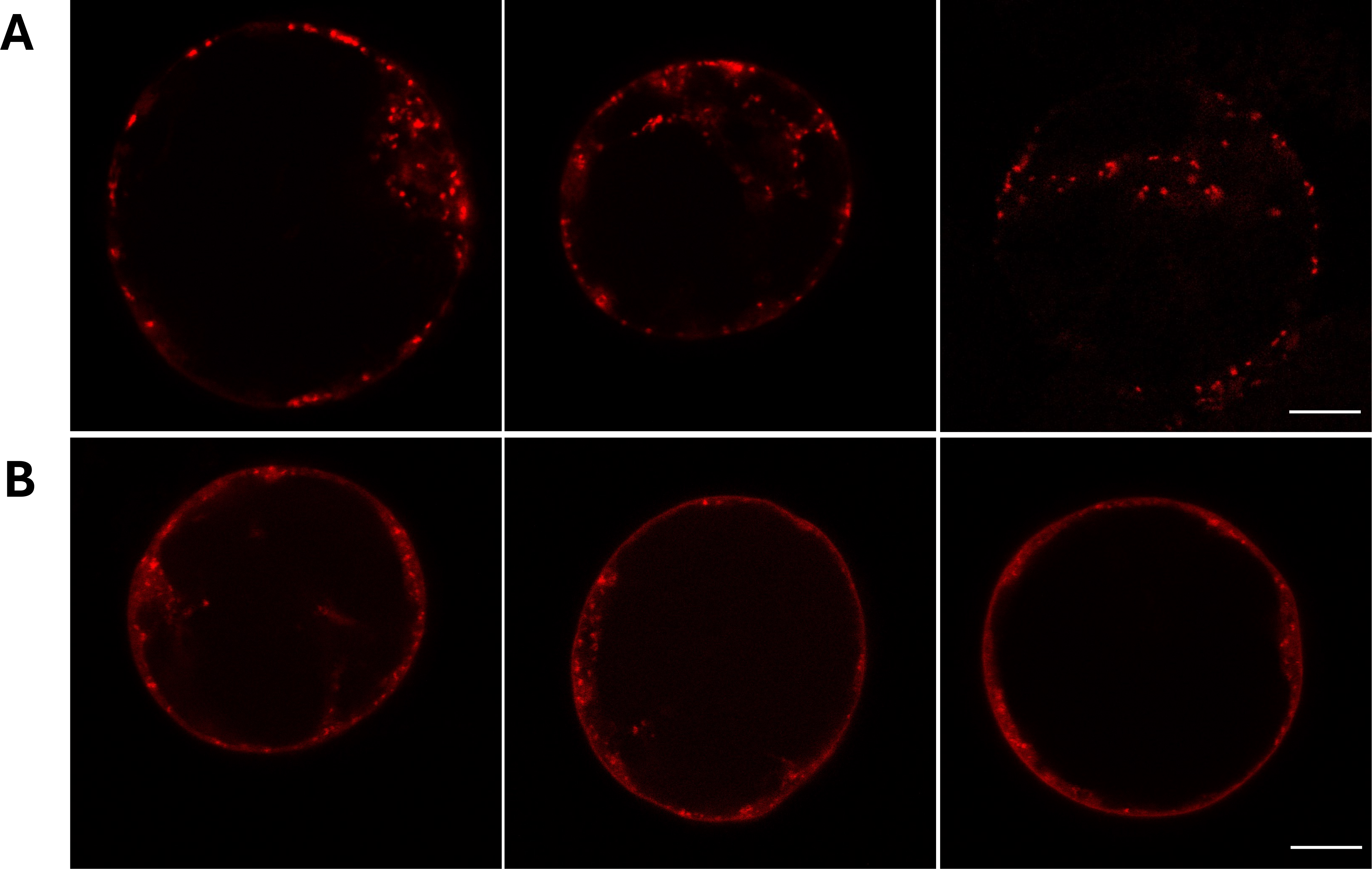

**Figure S5. Expression of the Goligi and TGN markers in PSBD cells.** *Arabidopsis* protoplasts were transfected with the Golgi (ManI-mRFP) (A) or TGN (B) markers before confocal imaging of transfected cells. Scale bar =10 μm.

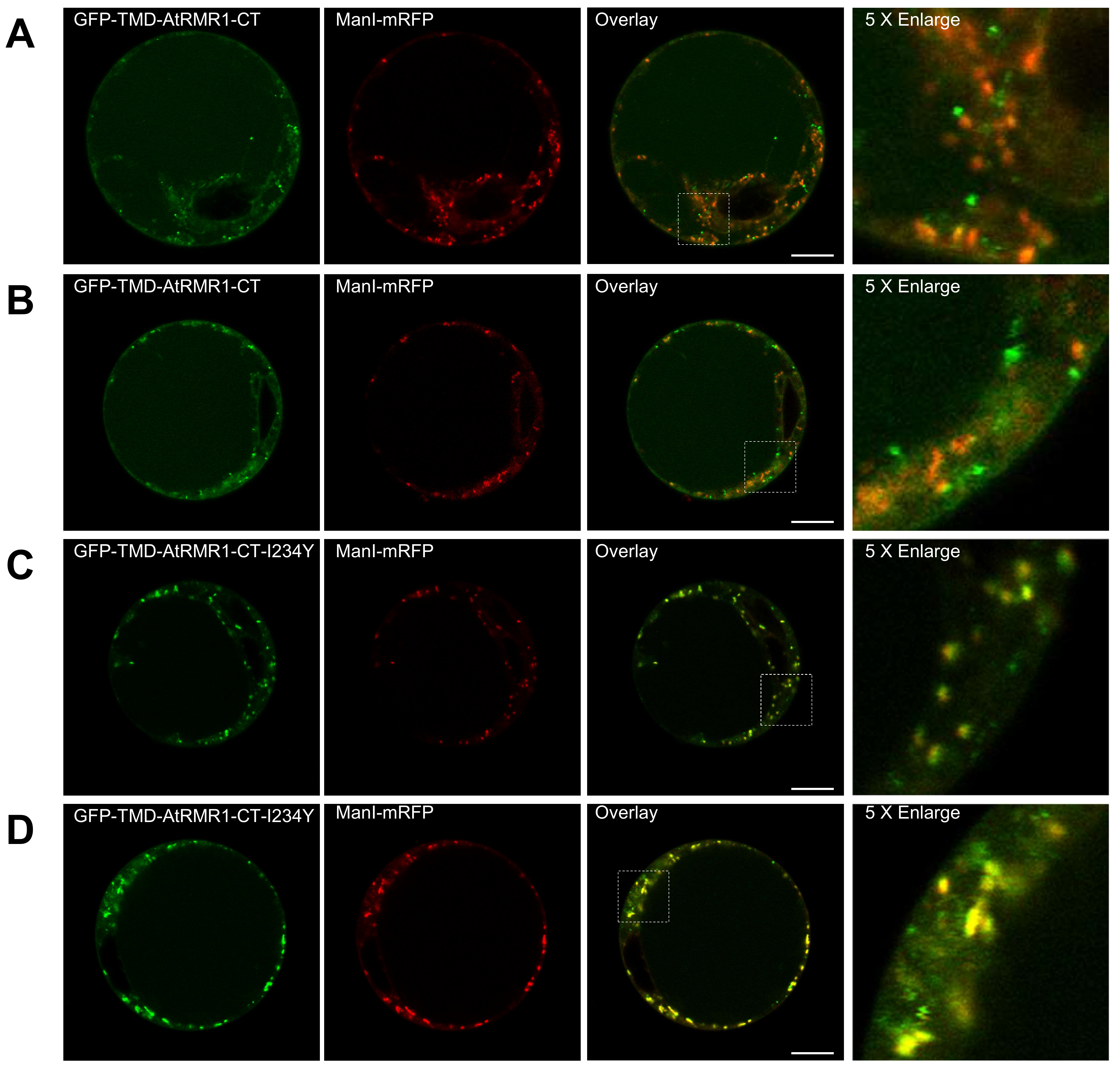

**Figure S6. Confocal images from additional biological repeats used in the colocalization analysis of GFP-TMD-AtRMR1-CT and the Golgi marker.** *Arabidopsis* protoplasts were transfected with (A and B) wild-type or (C and D) I234Y mutant of GFP-TMD-AtRMR1-CT and the Golgi marker (ManI-mRFP) before confocal imaging of transfected cells. (Scale bar, 10 μm).

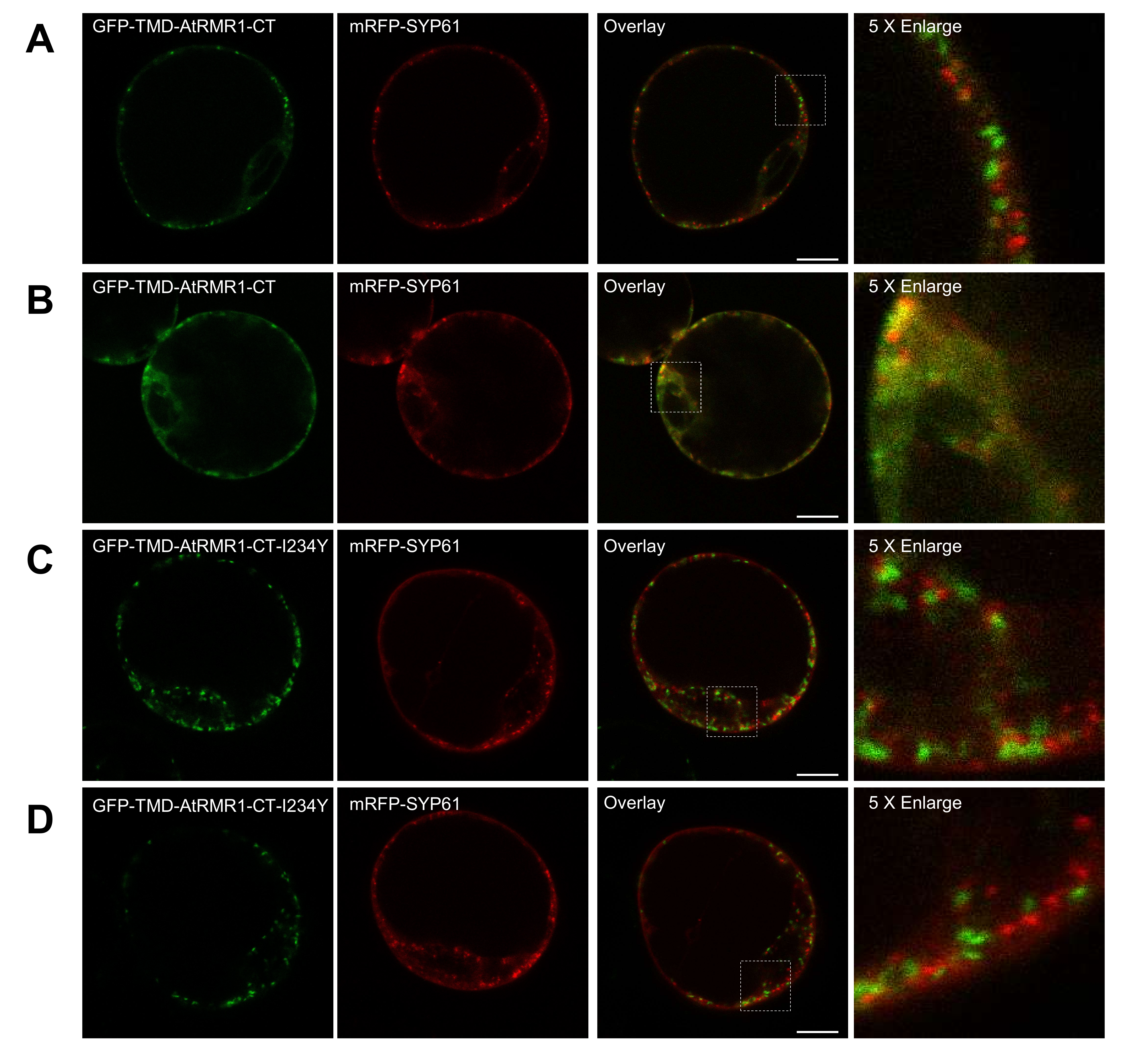

**Figure S7. Confocal images from additional biological repeats used in the colocalization analysis of GFP-TMD-AtRMR1-CT and the TGN marker.** *Arabidopsis* protoplasts were transfected with (A and B) wild-type or (C and D) I234Y mutant of GFP-TMD-AtRMR1-CT and the TGN marker (mRFP-SYP61) before confocal imaging of transfected cells. (Scale bar, 10 μm).

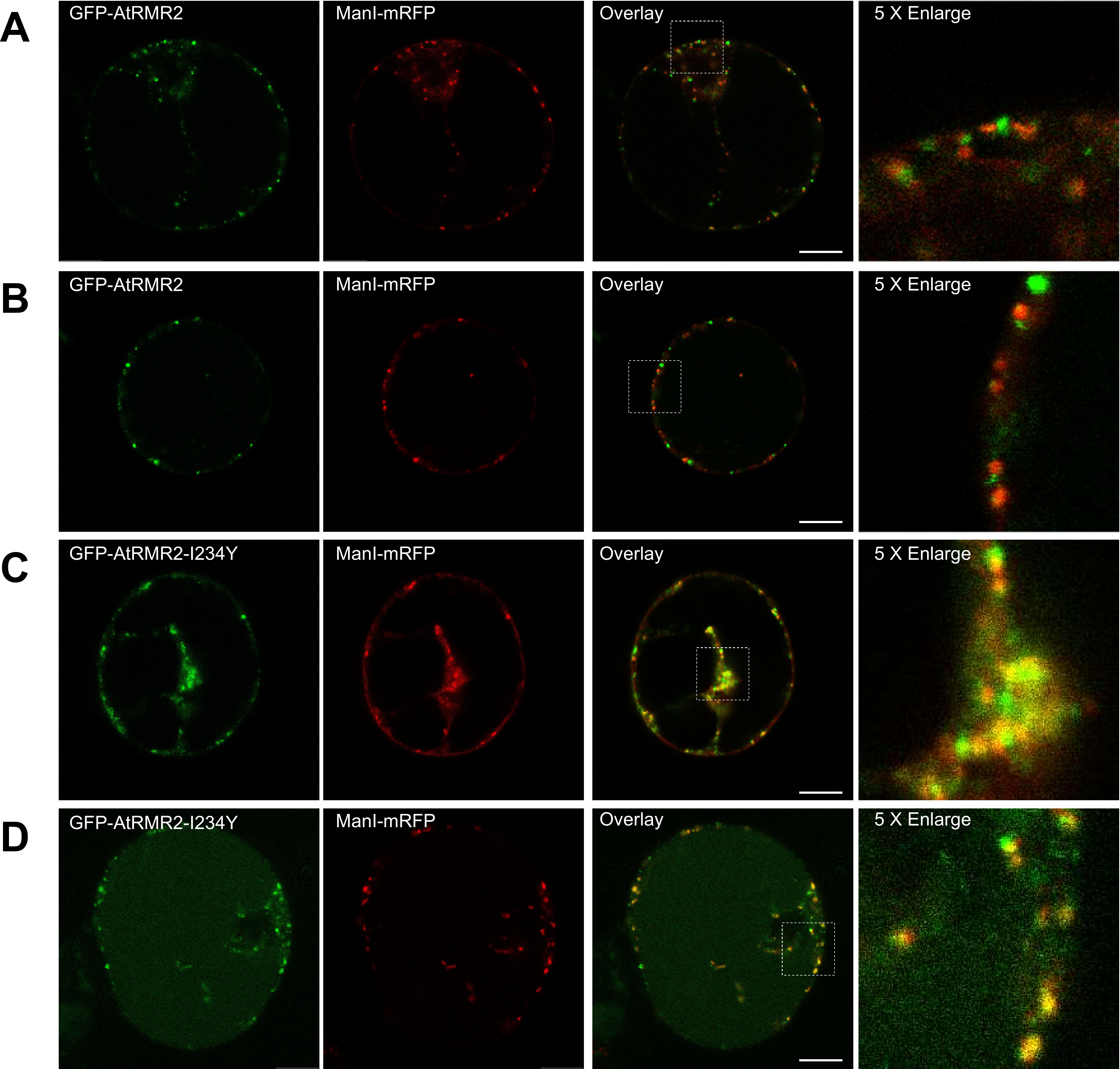

**Figure S8. Confocal images from additional biological repeats used in the colocalization analysis of GFP-AtRMR2 and the Golgi marker.** *Arabidopsis* protoplasts were transfected with (A and B) wild-type or (C and D) I234Y mutant of GFP-AtRMR2 with the Golgi marker (ManI-mRFP) before confocal imaging of transfected cells. (Scale bar, 10 μm).

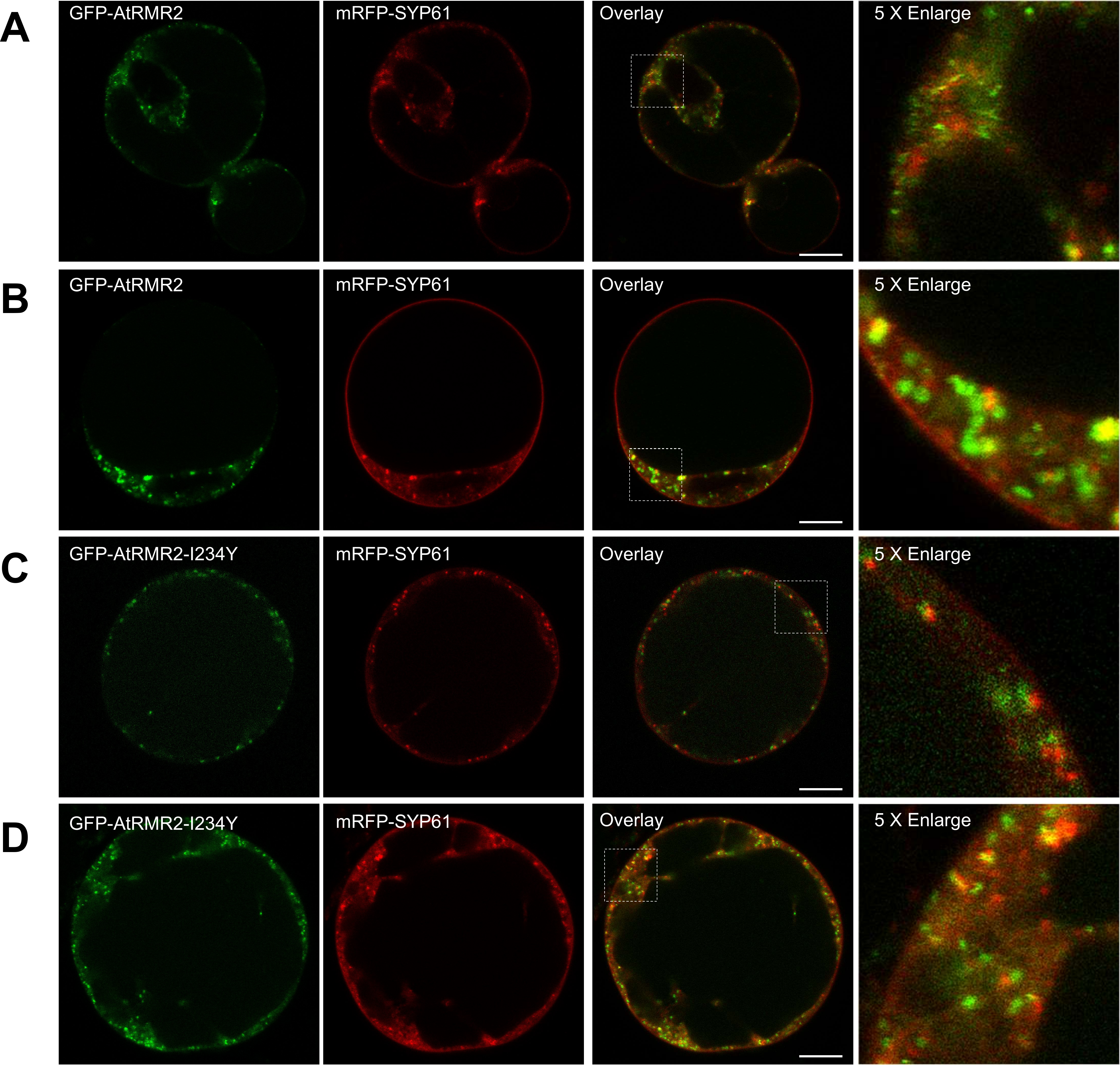

**Figure S9. Confocal images from additional biological repeats used in the colocalization analysis of GFP-AtRMR2 and the TGN marker.** *Arabidopsis* protoplasts were transfected with (A and B) wild-type or (C and D) I234Y mutant of GFP-AtRMR2 with the TGN marker (mRFP-SYP61) before confocal imaging of transfected cells. (Scale bar, 10 μm).

**Table S1. Data collection and refinement statistics of AtRMR1-RING**

**Data Collection**

Space group	C2

Cell dimensions

a, b, c (Å)	74.431, 23.648, 75.699

α, β, γ (°)	90, 94.8, 90

Resolution (Å)	37.72-2.10 (2.16-2.10)

No. of reflections	7849 (650)

R_merge_ (%)	8.5 (19.1)

I/σ (I)	9.0 (5.0)

Completeness	98.5 (97.4)

Multiplicity	3.6 (3.4)

**Refinement**

R_work_/R_free_ (%)	17.91/22.19

No. of atoms	1,263

Protein	1,184

Water	73

Metal	6

r.m.s. deviation

Bond lengths (Å)	0.007

Bond angles (°)	0.478

Clashscore	1.8

Ramachandran Plot

Favored	98%

Allowed	2%

Outliers	0%

**Table S2.** **Protein and Nucleotide sequences used in this study**

| **Constructs** | **Protein sequence** | **Nucleotide sequence** |
| --- | --- | --- |
| AtUba1  (72-1080) | EIDEDLHSRQLAVYGRETMRRLFASNVLISGMHGLGAEIAKNLILAGVKSVTLHDERVVELWDLSSNFVFSEDDVGKNRADASVQKLQDLNNAVVVSSLTKSLNKEDLSGFQVVVFSDISMERAIEFDDYCHSHQPPIAFVKADVRGLFGSVFCDFGPEFAVLDVDGEEPHTGIIASISNENQAFISCVDDERLEFEDGDLVVFSEVEGMTELNDGKPRKIKSTRPYSFTLDEDTTNYGTYVKGGIVTQVKQPKLLNFKPLREALKDPGDFLFSDFSKFDRPPLLHLAFQALDHFKAEAGRFPVAGSEEDAQKLISIATAINTGQGDLKVENVDQKLLRHFSFGAKAVLNPMAAMFGGIVGQEVVKACSGKFHPLFQFFYFDSVESLPSEPVDSSDFAPRNSRYDAQISVFGAKFQKKLEDAKVFTVGSGALGCEFLKNLALMGVSCGSQGKLTVTDDDIIEKSNLSRQFLFRDWNIGQAKSTVAASAAAVINPRFNIEALQNRVGAETENVFDDAFWENLTVVVNALDNVNARLYVDSRCLYFQKPLLESGTLGTKCNTQSVIPHLTENYGASRDPPEKQAPMCTVHSFPHNIDHCLTWARSEFEGLLEKTPAEVNAYLSSPVEYTNSMMSAGDAQARDTLERIVECLEKEKCETFQDCLTWARLRFEDYFVNRVKQLIYTFPEDAATSTGAPFWSAPKRFPRPLQYSSSDPSLLNFITATAILRAETFGIPIPEWTKNPKEAAEAVDRVIVPDFEPRQDAKIVTDEKATTLTTASVDDAAVIDDLIAKIDQCRHNLSPDFRMKPIQFEKDDDTNYHMDVIAGLANMRARNYSIPEVDKLKAKFIAGRIIPAIATSTAMATGLVCLELYKVLDGGHKVEAYRNTFANLALPLFSMAEPLPPKVVKHRDMAWTVWDRWVLKGNPTLREVLQWLEDKGLSAYSISCGSCLLFNSMFTRHKERMDKKVVDLARDVAKVELPPYRNHLDVVVACEDEDDNDVDIPLVSIYFR | GAGATTGACGAAGATCTGCACAGCCGTCAACTGGCGGTTTACGGTCGTGAAACCATGCGTCGTCTGTTCGCGAGCAACGTGCTGATCAGCGGCATGCATGGTCTGGGCGCGGAAATCGCGAAAAACCTGATTCTGGCGGGTGTGAAGAGCGTTACCCTGCACGACGAGCGTGTGGTTGAACTGTGGGATCTGAGCAGCAACTTCGTGTTTAGCGAGGACGATGTTGGCAAAAACCGTGCGGACGCGAGCGTTCAGAAGCTGCAAGATCTGAACAACGCGGTGGTTGTGAGCAGCCTGACCAAAAGCCTGAACAAGGAAGACCTGAGCGGTTTCCAGGTTGTGGTTTTTAGCGATATCAGCATGGAGCGTGCGATTGAGTTCGACGATTACTGCCACAGCCACCAACCGCCGATCGCGTTCGTGAAGGCGGACGTTCGTGGTCTGTTTGGCAGCGTGTTCTGCGATTTTGGTCCGGAGTTCGCGGTGCTGGACGTTGATGGCGAGGAACCGCACACCGGCATCATTGCGAGCATCAGCAACGAGAACCAGGCGTTTATTAGCTGCGTTGACGATGAACGTCTGGAGTTCGAAGACGGTGATCTGGTGGTTTTTAGCGAGGTGGAGGGTATGACCGAGCTGAACGACGGCAAACCGCGTAAGATCAAAAGCACCCGTCCGTATAGCTTCACCCTGGACGAAGATACCACCAACTACGGCACCTATGTTAAGGGTGGCATTGTGACCCAGGTTAAACAACCGAAGCTGCTGAACTTTAAGCCGCTGCGTGAGGCGCTGAAGGACCCGGGTGATTTCCTGTTTAGCGACTTCAGCAAATTTGATCGTCCGCCGCTGCTGCACCTGGCGTTCCAGGCGCTGGACCACTTTAAAGCGGAAGCGGGTCGTTTCCCGGTGGCGGGCAGCGAGGAAGATGCGCAAAAGCTGATCAGCATTGCGACCGCGATCAACACCGGTCAGGGCGACCTGAAAGTGGAGAACGTTGATCAAAAGCTGCTGCGTCACTTCAGCTTTGGCGCGAAGGCGGTTCTGAACCCGATGGCGGCGATGTTTGGTGGCATTGTGGGTCAGGAAGTGGTTAAAGCGTGCAGCGGCAAGTTCCACCCGCTGTTTCAATTCTTTTACTTCGACAGCGTTGAGAGCCTGCCGAGCGAACCGGTGGACAGCAGCGATTTTGCGCCGCGTAACAGCCGTTATGACGCGCAGATCAGCGTTTTCGGTGCGAAATTTCAAAAGAAACTGGAGGATGCGAAGGTGTTCACCGTTGGTAGCGGCGCGCTGGGTTGCGAATTTCTGAAAAACCTGGCGCTGATGGGCGTTAGCTGCGGTAGCCAGGGCAAACTGACCGTGACCGACGATGACATCATTGAGAAGAGCAACCTGAGCCGTCAGTTCCTGTTTCGTGACTGGAACATCGGCCAAGCGAAGAGCACCGTGGCGGCGAGCGCGGCGGCGGTTATCAACCCGCGTTTCAACATTGAGGCGCTGCAAAACCGTGTGGGTGCGGAAACCGAAAACGTTTTCGATGACGCGTTTTGGGAAAACCTGACCGTGGTTGTGAACGCGCTGGACAACGTGAACGCGCGTCTGTACGTTGATAGCCGTTGCCTGTATTTTCAGAAACCGCTGCTGGAAAGCGGCACCCTGGGCACCAAGTGCAACACCCAAAGCGTTATCCCGCACCTGACCGAGAACTACGGTGCGAGCCGTGACCCGCCGGAAAAACAGGCGCCGATGTGCACCGTGCACAGCTTCCCGCACAACATTGATCACTGCCTGACCTGGGCGCGTAGCGAGTTTGAAGGCCTGCTGGAGAAGACCCCGGCGGAAGTTAACGCGTACCTGAGCAGCCCGGTGGAGTATACCAACAGCATGATGAGCGCGGGTGATGCGCAGGCGCGTGATACCCTGGAGCGTATCGTGGAATGCCTGGAGAAGGAAAAATGCGAAACCTTTCAAGATTGTTTAACCTGGGCGCGTCTGCGTTTCGAAGATTACTTTGTGAACCGTGTTAAACAACTGATTTATACCTTTCCGGAAGATGCGGCGACCAGCACCGGTGCGCCGTTCTGGAGCGCGCCGAAGCGTTTTCCGCGTCCGCTGCAGTACAGCAGCAGCGACCCGAGCCTGCTGAACTTCATCACCGCGACCGCGATTCTGCGTGCGGAAACCTTTGGTATCCCGATTCCGGAATGGACCAAAAACCCGAAAGAGGCGGCGGAAGCGGTTGACCGTGTGATCGTTCCGGATTTCGAGCCGCGTCAAGACGCGAAAATTGTGACCGATGAAAAAGCGACCACCCTGACCACCGCGAGCGTGGATGATGCGGCGGTTATCGATGACCTGATCGCGAAAATTGACCAGTGCCGTCACAACCTGAGCCCGGATTTCCGTATGAAACCGATTCAATTTGAGAAGGATGACGATACCAACTACCACATGGACGTTATCGCGGGTCTGGCGAACATGCGTGCGCGTAACTATAGCATTCCGGAAGTGGATAAGCTGAAAGCGAAGTTCATCGCGGGTCGTATCATTCCGGCGATTGCGACCAGCACCGCGATGGCGACCGGCCTGGTGTGCCTGGAGCTGTACAAAGTTCTGGACGGTGGCCACAAGGTGGAAGCGTATCGTAACACCTTCGCGAACCTGGCGCTGCCGCTGTTTAGCATGGCGGAGCCGCTGCCGCCGAAAGTTGTGAAGCACCGTGACATGGCGTGGACCGTGTGGGATCGTTGGGTTCTGAAAGGTAACCCGACCCTGCGTGAGGTTCTGCAGTGGCTGGAAGACAAGGGTCTGAGCGCGTACAGCATCAGCTGCGGCAGCTGCCTGCTGTTCAACAGCATGTTTACCCGTCACAAAGAGCGTATGGATAAGAAAGTTGTGGACCTGGCGCGTGATGTGGCGAAGGTTGAACTGCCGCCGTACCGTAACCACCTGGACGTTGTGGTTGCGTGCGAGGATGAAGACGATAACGACGTGGATATCCCGCTGGTTAGCATTTATTTCCGT |
| AtUbc30  (1-148) | MASKRINKELRDLQRDPPVSCSAGPTGDDMFQWQATIMGPADSPFAGGVFLVTIHFPPDYPFKPPKVAFRTKVYHPNINSNGSICLDILKEQWSPALTVSKVLLSICSLLTDPNPDDPLVPEIAHIYKTDRVKYESTAQSWTQKYAMG | ATGGCGAGCAAGCGTATCAACAAAGAGCTGCGTGACCTGCAGCGTGATCCGCCGGTTAGCTGCAGCGCGGGTCCGACCGGCGACGATATGTTCCAGTGGCAAGCGACCATCATGGGTCCGGCGGACAGCCCGTTTGCGGGTGGCGTGTTTCTGGTTACCATTCACTTCCCGCCGGATTACCCGTTTAAGCCGCCGAAAGTGGCGTTTCGTACCAAGGTTTATCACCCGAACATCAACAGCAACGGCAGCATCTGCCTGGACATTCTGAAGGAACAATGGAGCCCGGCGCTGACCGTGAGCAAAGTTCTGCTGAGCATTTGCAGCCTGCTGACCGACCCGAACCCGGATGATCCGCTGGTGCCGGAGATCGCGCACATTTACAAGACCGATCGTGTTAAATATGAAAGCACCGCGCAGAGCTGGACCCAAAAATACGCGATGGGT |
| AtUb  (1-76) | MQIFVKTLTGKTITLEVESSDTIDNVKAKIQDKEGIPPDQQRLIFAGKQLEDGRTLADYNIQKESTLHLVLRLRGG | ATGCAGATTTTTGTGAAGACCCTGACCGGCAAGACCATTACCCTGGAAGTTGAAAGCAGCGACACCATTGATAACGTGAAAGCGAAAATCCAGGACAAAGAGGGTATCCCGCCGGATCAGCAACGTCTGATCTTCGCGGGTAAACAACTGGAAGACGGCCGTACCCTGGCGGATTACAACATTCAAAAGGAGAGCACCCTGCATCTGGTTCTGCGTCTGCGTGGCGGT |
| AtRMR1-RING (206-283) | RLDAKLVHTLPCFTFTDSAHHKAGETCAICLEDYRFGESLRLLPCQHAFHLNCIDSWLTKWGTSCPVCKHDIRTETMS | CGTCTGGACGCGAAGCTGGTGCACACCCTGCCGTGCTTCACCTTTACCGATAGCGCGCACCACAAAGCGGGTGAAACCTGCGCGATTTGCCTGGAAGACTACCGTTTTGGTGAAAGCCTGCGTCTGCTGCCGTGCCAACACGCGTTTCACCTGAACTGCATCGACAGCTGGCTGACCAAGTGGGGTACCAGCTGCCCGGTGTGCAAACACGATATTCGTACCGAAACCATGAGC |
| AtRMR2-RING (207-283) | SRRLVKAMPSLIFSSFHEDNTTAFTCAICLEDYTVGDKLRLLPCCHKFHAACVDSWLTSWRTFCPVCKRDARTSTGE | AGCCGTCGTCTGGTTAAGGCGATGCCGAGCCTGATCTTCAGCAGCTTTCACGAGGACAACACCACCGCGTTTACCTGCGCGATTTGCCTGGAAGACTACACCGTTGGCGATAAGCTGCGTCTGCTGCCGTGCTGCCACAAATTTCATGCGGCGTGCGTGGACAGCTGGCTGACCAGCTGGCGTACCTTTTGCCCGGTTTGCAAACGTGATGCGCGTACCAGCACCGGCGAG |
| AtRMR3-RING (207-282) | CRRTVKAMPSVTFTCAKIDNTTGFSCAICLEDYIVGDKLRVLPCSHKFHVACVDSWLISWRTFCPVCKRDARTTAD | TGCCGTCGTACCGTGAAGGCGATGCCGAGCGTTACCTTCACCTGCGCGAAAATTGACAACACCACCGGTTTTAGCTGCGCGATCTGCCTGGAGGACTATATTGTGGGCGATAAGCTGCGTGTTCTGCCGTGCAGCCACAAATTCCACGTGGCGTGCGTTGATAGCTGGCTGATCAGCTGGCGTACCTTTTGCCCGGTGTGCAAACGTGATGCGCGTACCACCGCGGAT |
| AtRMR4-RING (209-284) | PKSMIIRMPTTIFNGICDEATTSILCCICLENYEKGDKLRILPCHHKFHVACVDLWLGQRKSFCPVCKRDARSIST | CCGAAGAGCATGATCATTCGTATGCCGACCACCATCTTTAACGGCATTTGCGATGAGGCGACCACCAGCATCCTGTGCTGCATTTGCCTGGAGAACTACGAAAAGGGTGACAAACTGCGTATCCTGCCGTGCCACCACAAGTTCCACGTGGCGTGCGTTGATCTGTGGCTGGGCCAGCGTAAGAGCTTTTGCCCGGTGTGCAAACGTGACGCGCGTAGCATTAGCACC |

**Table S3.** **Constructs and primers used in this study**

| Constructs / Uses | Primer sequence (5’ to 3’) |
| --- | --- |
| **pGEX-6p-1-AtRMR1-RING**  Subcloning of AtRMR1-RING (206-283) into the vector pGEX-6p-1 for E. coli expression. | CGCGGATCCCGTCTGGACGCGAAGCTG |
|  | ACGCGTCGACTCAGCTCATGGTTTCGGTACG |
| **pGEX-6p-1-AtUba1**  Subcloning of AtUba1 (72-1080) into the vector pGEX-6p-1 for E. coli expression. | CGCGGATCCGAGATTAGACGAAGATCTG |
|  | ACGCGTCGACTCAACGGAAATAAATGCTAAC |
| **pGEX-6p-1-AtUbc30**  Subcloning of AtUbc30 into the vector pGEX-6p-1 for E. coli expression. | CGCGGATCCATGGCGAGCAAGCGTATC |
|  | ACGCGTCGACTCAGCCCATCGCGTATT |
| **pGEX-6p-1-AtUb**  Subcloning of AtUb into the vector pGEX-6p-1 for E. coli expression. | CGCGGATCCATGCAGATTTTTGTG |
|  | ACGCGTCGACTCAACCGCCACGCAG |
| **GFP-TMD-AtRMR1-CT**  Subcloning of TMD-AtRMR1-CT (166-310) into the vector pBI221-AluSP-GFP for transient expression in Arabidopsis protoplasts | CTAGTCTAGAGCTTGGACTGTGTTG |
|  | CGAGCTCCTAACGGCTTTGACTG |
| **GFP-AtRMR2**  Subcloning of AtRMR2 (22-448) into the vector pBI221-AluSP-GFP for transient expression in Arabidopsis protoplasts | CTAGTCTAGAGTTATTTTGATGAGGAATAACATC |
|  | CGAGCTCCTAACAGTCTGGAAGCGAG |
| **AtRMR1-RING-I234Y**  To introduce I234Y mutation in pGEX-6p-1-AtRMR1-RING | AACCTGCGCGTACTGCCTGGAAGAC |
|  | TCACCCGCTTTGTGGTGC |
| **GFP-TMD-AtRMR1-CT-I234Y**  To introduce I234Y mutation in GFP-TM-AtRMR1-CT | AACATGTGCTTACTGTCTCGAGGATTACAGATTTGGAGAAAGCCTCAG |
|  | TCCCCGGCCTTGTGGTGA |
